# Impact of Prior Infection on Protection and Transmission of SARS-CoV-2 in Golden Hamsters

**DOI:** 10.1101/2021.01.30.428920

**Authors:** Cheng Zhang, Zhendong Guo, Nan Li, Huan Cui, Keyin Meng, Lina Liu, Li Zhao, Shanshan Zhang, Chengfeng Qin, Juxiang Liu, Yuwei Gao, Chunmao Zhang

**Author notes:** Authors contributed equally to this work (Cheng Zhang, Zhendong Guo, Nan Li). Correspondence author: Chunmao Zhang, Yuwei Gao.

## Abstract

The severe acute respiratory syndrome coronavirus 2 (SARS-CoV-2) has caused over 100 million confirmed human infections, and 2 million more deaths globally since its emergence in the end of 2019. Several studies have shown that prior infection provided protective immunity against SARS-CoV-2 in non-human primate models. However, the effect of prior infection on blocking SARS-CoV-2 transmission is not clear. Here, we evaluated the impact of prior infection on protection and transmission of the SARS-CoV-2 virus in golden hamsters. Our results showed that prior infection provided protective immunity against SARS-CoV-2 re-challenge, but it was not sterizing immunity. The transmission experiment results showed that SARS-CoV-2 was efficiently transmitted from naive hamsters to prior infected hamsters by direct contact and airborne route, but not by indirect contact. Further, the virus was efficiently transmitted from prior infected hamsters to naive hamsters by direct contact, but not by airborne route and indirect contact. Surprisingly, the virus can be transmitted between prior infected hamsters by direct contact during a short period of early infection. Taken together, our study demonstrated that prior infected hamsters with good immunity can still be naturally re-infected, and the virus can be transmitted between prior infected hamsters and the naive through different transmission routes, implying the potential possibility of human re-infection and the risk of virus transmission between prior infected population and the healthy. Our study will help to calculate the herd immunity threshold more accurately, make more reasonable public health decisions, formulate an optimized population vaccination program, as well as aid the implementation of appropriate public health and social measures to control COVID-19.

## The main text

As of Jan 29, 2021, more than 100 million confirmed human infections and over 2 million deaths have been caused by the severe acute respiratory syndrome coronavirus 2 (SARS-CoV-2), the causative agent of coronavirus infectious disease 2019 (COVID-19) pandemic, with devastating impact on lives and economy. At present, multiple vaccine candidates are in phase 3 clinical trials and several of them have been approved for emergency use authorization with conditions^1–8^. Previous studies have shown that prior infection or vaccination provided protective immunity against SARS-CoV-2 in different animal models^9–17^, but most are not sterizing immunity. The prior infected or vaccinated animals still shed large quantities of virus in their upper respiratory tracts^9,12,14,16^. Besides protection from diseases, reducing or blocking SARS-CoV-2 transmission between humans is crucial for COVID-19 pandemic control. However, the impact of prior infection on blocking SARS-CoV-2 transmission is not clear. As a small animal model, golden hamsters have been used for studying pathogenesis^18,19^, transmission ability^20^ of SARS-CoV-2 and for evaluating potential vaccines^21,22^ and antiviral drugs^23^. Here, we evaluated the impact of prior infection on protection and transmission of SARS-CoV-2 in golden hamsters. Our results showed that prior infection provided good protective immunity against SARS-CoV-2 re-challenge, but the prior infected hamsters can still be re-infected. Moreover, the virus was efficiently transmitted from naive hamsters to prior infected hamsters by direct contact and airborne route, but not by indirect contact. The virus was also transmitted from prior infected hamsters to the naive and prior infected hamsters by direct contact, but not by airborne route. Our findings will help governments and public health agencies to make more reasonable public health decisions as well as aid the implementation of appropriate public health and social measures to control COVID-19.

### Prior infection protects hamsters against SARS-CoV-2 re-challenge

We evaluated protective immunity of prior infection against SARS-CoV-2 re-challenge. Hamsters were divided into high dose infected group (HD) and low dose infected group (LD), and intranasally inoculated with 10^5^ TCID_50_ or 10^3^ TCID_50_ of the virus respectively (Supplementary Table S1). At 21 days post infection (dpi), Hamsters in HD and LD were inoculated with 10^6^ TCID_50_ of the virus. As the infected control (IC), another six naive hamsters were inoculated with 10^6^ TCID_50_ of the virus. At 2 and 4 dpi, nasal washes and the supernatants of the homogenized nasal turbinates and lungs were collected for virus titration in Vero-E6 cells and RNA quantification using real-time qPCR. The glutaraldehyde-fixed nasal turbinates were prepared for histological examination.

Serum was collected from hamsters in HD and LD at 21 dpi. The results of virus neutralization assay revealed that all hamsters inoculated with the virus had a much higher neutralizing antibody titer, and the neutralizing antibody titer in HD was slightly higher than that in LD (Figure S1), indicating that prior infection elicited effective immunity in hamsters. For nasal washes, viral load in IC peaked at 2 dpi, with a titer of 10^5.17^ TCID_50_/mL, and was significantly much higher than that in HD and LD (Figure 1A). At 2 and 4 dpi, RNA copies in IC were slightly higher than that in HD and LD with significant difference (Figure 1D). For nasal turbinate, high levels of viral load were observed in IC at 2 and 4 dpi, with a titer of 10^5.75^ TCID_50_/mL and 10^3.75^ TCID_50_/mL respectively, about 3500-fold and 180-fold higher than that in HD and LD (Figure 1B). Viral load in HD and LD decreased to below the detection limit of TCID_50_ assay at 4 dpi. RNA copies in IC at 2 dpi, 10^10.45^copies/mL, were 80-fold and 25-fold higher than that in HD and LD respectively and RNA copies in IC at 4 dpi, 10^9.83^copies/mL, were about 4000-fold and 10000-fold higher than that in HD and LD (Figure 1E). Compared with at 2 dpi, RNA copies at 4 dpi in HD and LD decreased about 200-fold and 1600-fold respectively (Figure 1E). Viral RNA assays were further confirmed by the sgmRNA assays. The sgmRNA copies in IC at 2 and 4 dpi, 10^7.1^copies/mL and 10^6.4^copies/mL, were about 80-fold and 5000-fold higher than that in HD and LD, averagely 10^5.2^copies/mL and 10^2.7^copies/mL respectively (Figure 1G). Compared with at 2 dpi, the sgmRNA copies in HD and LD at 4 dpi decreased about 250-fold and 600-fold respectively (Figure 1G). For lungs, viral load at 2 and 4 dpi in HD and LD was under the detection limit, which was significantly lower than that in IC, 10^5.33^ TCID_50_/mL and 10^4.17^ TCID_50_/mL respectively (Figure 1C). RNA copies at 2 and 4 dpi in IC, 10^10.2^copies/mL and 10^9.5^copies/mL, was about 13000-fold and 40000-fold higher than that in HD and LD, about 10^6.1^copies/mL and 10^4.9^copies/mL (Figure 1F). Similar to the trend of viral RNA copies, the sgmRNA in IG at 2 and 4 dpi, 10^6.6^copies/mL and 10^6.0^copies/mL, was about 16000-fold and 8000-fold higher than that in HD and LD, averagely 10^2.4^copies/mL and 10^2.1^copies/mL respectively (Figure 1H). Additionally, we examined the presence of SARS-CoV-2 virus in nasal tissues using transmission electron microscopy. Several coronavirus-like particles were observed in intracellular compartments of nasal tissues of hamsters that were re-challenged with SARS-CoV-2 (Figure S2).

**Figure 1.**
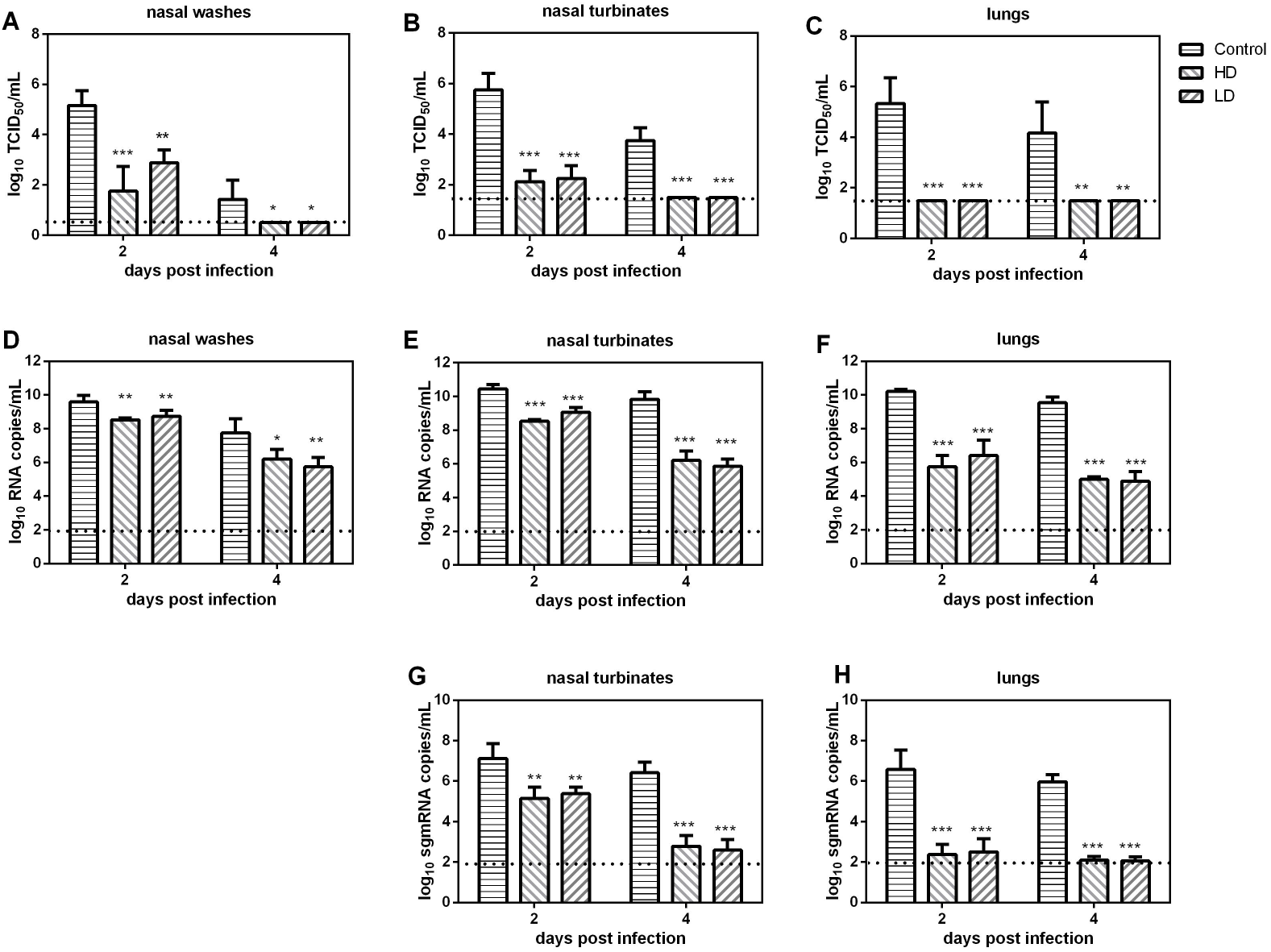
Viral load and histological examination in prior infected hamsters intranasally inoculated with SARS-CoV-2. Sixteen hamsters were randomly divided into HD and LD groups and inoculated with 10^5^ TCID_50_ or 10^3^ TCID_50_ of the SARS-CoV-2 virus respectively. At 21 dpi, hamsters in HD and LD were re-challenged with 10^6^ TCID_50_ of the SARS-CoV-2 virus. At 2 and 4 dpi, nasal washes, nasal turbinate and lungs were collected from hamsters for viral titration, RNA quantification and transmission electron microscopy examination. (A to C) Viral titers (log_10_TCID_50_/mL) detected in nasal washes (A), nasal turbinates (B) and lungs (C) of prior infected hamsters challenged with the SARS-CoV-2 virus. (D to F) Viral RNA copies (log_10_RNA copies/mL) detected in nasal washes (D), nasal turbinates (E) and lungs (F) of prior infected hamsters challenged with SARS-CoV-2. (G and H) Viral sgmRNA copies (log_10_sgmRNA copies/mL) detected in nasal turbinates (G) and lungs (H) of prior infected hamsters re-challenged with SARS-CoV-2. One-way analysis of variance (ANOVA) and Tukey’s multiple comparisons test were used to analyze the statistical differences of viral titers, RNA copies and sgmRNA copies in nasal washes, nasal turbinates and lungs between different experimental groups (p > 0.05, not significant, [ns]; p < 0.05, * p <0.01, **; p < 0.001, ***).

The substantially reduced viral titers, RNA and sgmRNA copies in nasal washes, nasal turbinate and lungs showed that prior infection provided good protective immunity against SARS-CoV-2. However, a moderate level of live virus was still detected in nasal washes and nasal turbinates, despite with a relatively short shedding period, and considerable sgmRNA copies were also detected in nasal turbinate at 2 dpi. The results of the detected live virus and the considerable sgmRNA copies, in combined with observation of coronavirus-like particles in intracellular compartments of nasal cells, powerfully proved that the virus can replicate in prior infected hamsters, especially in nasal turbinates, undoubtedly indicating that prior infected hamsters can be re-infected by the virus, even with a higher neutralizing antibody titer.

### Impact of prior infection on SARS-CoV-2 transmission in hamsters

The SARS-CoV-2 virus was transmitted between hamsters via multiple routes, including direct contact, indirect contact and airborne transmission. Here we systematically evaluated the impact of prior infection on SARS-CoV-2 transmission between prior infected hamsters and the naive hamsters.

### Transmission of SARS-CoV-2 from naive hamsters to prior infected hamsters

For the potential transmission of the virus from naive hamsters to prior infected hamsters by direct contact, three naive donor hamsters were intranasally inoculated with 10^6^ TCID_50_ of the virus. After 24 hours’ inoculation, the three donors were transferred to a direct contact transmission cage (supplementary Figure S3A) and co-housed with another three prior infected hamsters that were inoculated with 10^5^ TCID_50_ or 10^3^ TCID_50_ of the virus 21 days ago. Nasal washes were collected every other day from the donors and the contacts for 8 days. For donors, the infectious viral load in nasal washes peaked at 2 dpi and then declined rapidly, while viral RNA copies was relatively stable during the first six infection days, and then substantially declined at 8 dpi (Figure 2A). At 1 days post exposure (dpe), live SARS-CoV-2 virus was detected in nasal washes of two prior infected contact hamsters, and one with a very low viral titer. At 3 dpe, live virus was detected in all three prior infected contact hamsters (Figure 2A). The viral titers in the contacts were much lower than that in the donor hamsters. Viral RNA copies in two prior infected contact hamsters were significantly improved at 3 dpe (Figure 2A). The experiment results showed that the virus was efficiently transmitted from the naïve donors to the prior infected contacts. For airborne transmission of the virus from naive hamsters to prior infected hamsters, three naive donor hamsters were inoculated with 10^6^ TCID_50_ of the virus, and at 24 hours’ inoculation, the donor hamsters and another three prior infected recipient hamsters were transferred to the airborne transmission cage, with two wire-mesh partition that prevented the direct and indirect contact between animals and allowed the spread of the virus through air (supplementary Figure S3B). At 1 dpe, live SARS-CoV-2 virus was detected in nasal washes of two prior infected recipient hamsters and one with a very low viral titer (Figure 2B). At 3 dpe, all three prior infected recipient hamsters were infected by SARS-CoV-2 and viral titers in nasal washes were significantly improved, and at 5 dpe, the viral titer in one prior infected recipient hamster was still relatively high, about 10^3.25^ TCID_50_/mL (Figure 2B). Viral RNA copies in nasal washes in the prior infected recipient hamsters peaked at 3 dpe, and one hamster still had a higher viral RNA copies in nasal washes at 5 dpe (Figure 2B). Therefore, the virus was efficiently transmitted from the naive to prior infected hamsters by airborne route as well. For indirect contact transmission of the virus from the naive to prior infected hamsters, three naive donor hamsters were inoculated with 10^6^ TCID_50_ of the virus, and at 48 hours’ inoculation, the donor hamsters were removed and transferred to a new cage, and another three prior infected recipient hamsters were placed into the initial cage housing the donor hamsters. During the whole experiment period, no live virus was detected in nasal washes of the three prior infected recipient hamsters (Figure 2C). The virus was not transmitted from the naive to prior infected hamsters by indirect contact. In summary, SARS-CoV-2 was efficiently transmitted from the naive donors to prior infected hamsters by direct contact and airborne transmission, but not by indirect contact.

**Figure 2.**
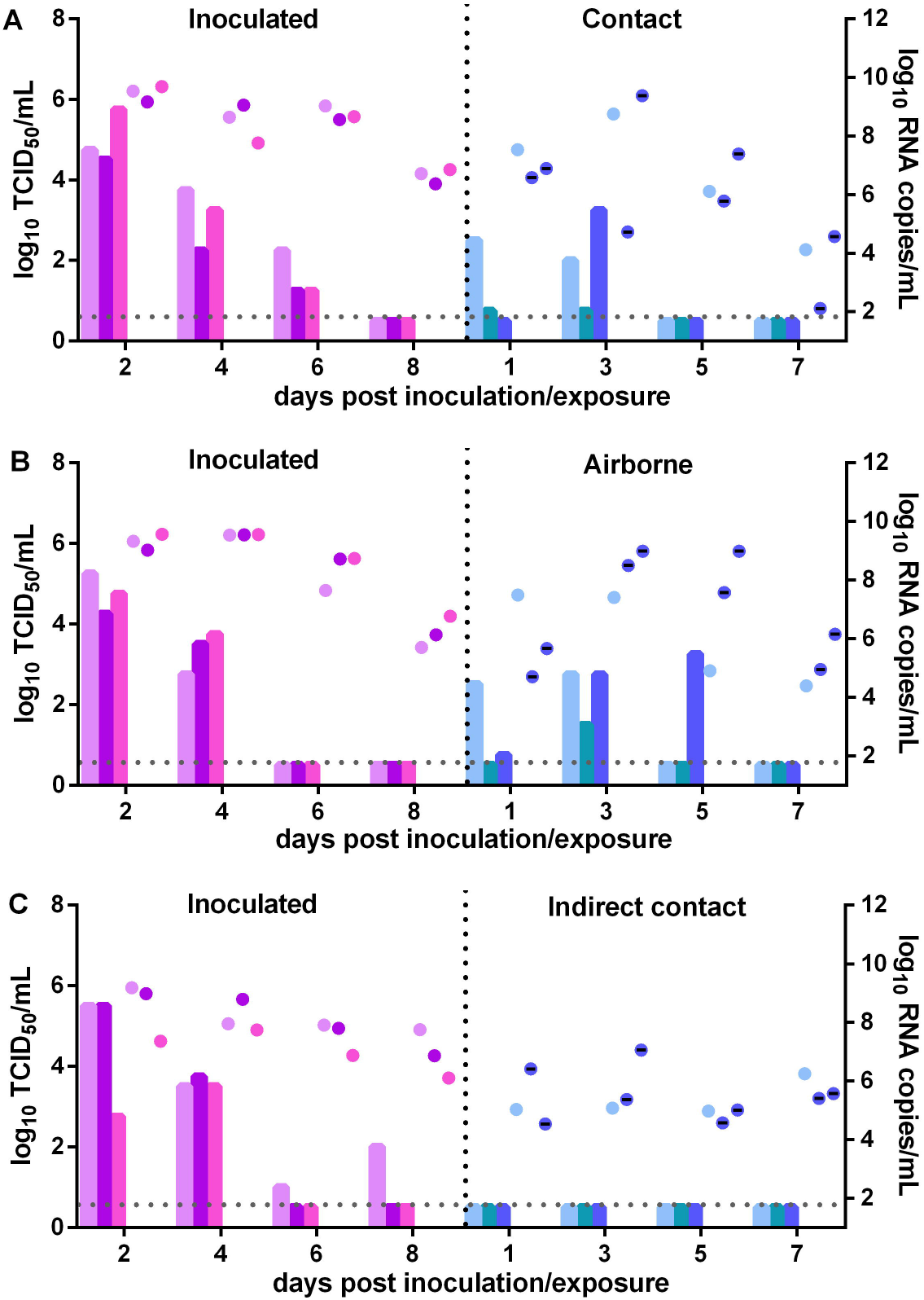
Transmission of SARS-CoV-2 from naïve hamsters to prior infected hamsters. (A) Infectious viral load (log_10_TCID_50_ shown in bars) and viral RNA copies (log_10_RNA copies/mL, shown in dots with matched color) detected in nasal washes of the naïve donor hamsters inoculated with 10^6^ TCID_50_ of SARS-CoV-2 and prior infected contact hamsters in the transmission group, which were previously inoculated with 10^5^ TCID_50_ or 10^3^ TCID_50_ of SARS-CoV-2 twenty-one days ago. At 24 hours’ inoculation, the donor hamsters and the prior infected contact hamsters were co-housed together in a new cage. (B) Viral titers and viral RNA copies detected in nasal washes of the naïve donor hamsters inoculated with SARS-CoV-2 and those prior infected hamsters in airborne transmission group. At 24 hours’ inoculation, the donor hamsters and the prior infected hamsters were transferred to an airborne transmission cage. (C) Viral titers and viral RNA copies detected in nasal washes of the naïve donor hamsters inoculated with SARS-CoV-2 and the prior infected hamsters in the indirect contact transmission group. At 48 hours’ inoculation, the donor hamsters were removed and transferred to a new cage and prior infected hamsters were placed into the cage housing the naïve hamsters before. Nasal washes were collected from all hamsters in different experiment groups every other day for virus titration and RNA quantification.

### Transmission of SARS-CoV-2 from prior infected hamsters to naive hamsters

For direct contact transmission of the virus from prior infected hamsters to naive hamsters, three prior infected hamsters were as the donors, and another three naive hamsters were as the recipients in direct contact transmission group. The donors were inoculated with 10^6^ TCID_50_ of the virus, and at 24 hours’ inoculation, the donors and the direct contacts were co-housed together in a new cage. At 2 dpi, viral titers in nasal washes of two donor hamsters were very low, and another donor hamster with a moderate titer of 10^2.75^ TCID_50_/mL (Figure 3A). At 4 dpi, live virus was not detected in nasal washes of two donor hamsters, and another one with a low viral titer. Viral RNA copies in nasal washes of the donor hamsters were about 10^8^copies/mL at 2 dpi, and substantially declined at 6 and 8 dpi (Figure 3A). Live virus was detected in nasal washes of all three contact hamsters at 3 days post exposure (dpe), one with a very high titer of 10^5.5^ TCID_50_/mL, and another with a very low titer of 10^0.75^ TCID_50_/mL (Figure 3A). Viral titers in nasal washes of all hamsters were very high at 5 dpe. Viral RNA copies in nasal washes of the contact hamsters were improved rapidly at 3 dpe and later held at a high level. The results showed that the virus was efficiently transmitted from prior infected hamsters to naive hamsters by direct contact. For airborne transmission of the virus from prior infected hamsters to naive hamsters, three prior infected donor hamsters were inoculated with 10^6^ TCID_50_ of the virus, and another three naive hamsters were as recipients. At 24 hours’ inoculation, the prior infected donor hamsters and the naïve recipient hamsters were transferred to an airborne transmission cage. Live virus was not detected at 2 dpi and later in nasal washes of the prior infected donor hamsters (Figure 3B). No live virus was also detected in nasal washes of the recipient hamsters in the airborne transmission group (Figure 3B). The airborne transmission experiment was also performed similarly at two hours’ inoculation. A moderate level of virus titer was detected at 1and 3 dpi in prior infected donor hamsters, and no live virus was still detected in all recipient hamsters in the airborne transmission group during the whole experiment period (Figure S4B). The results demonstrated that the virus was not transmitted from prior infected donors to naive hamsters through airborne route. For indirect contact transmission of the virus from prior infected hamsters to naive hamsters, three prior infected donor hamsters were inoculated with 10^6^ TCID_50_ of the virus, and at 48 hours’ inoculation, the prior infected donor hamsters were removed and transferred to a new cage, and another three naive recipient hamsters were placed into the initial cage housing the donors. No live virus was detected in nasal washes of the recipient hamsters in the indirect contact transmission group (Figure 3C). The virus was not transmitted from prior infected hamsters to the naive by indirect contact. To sum up, SARS-CoV-2 can be transmitted from prior infected hamsters to naive hamsters by direct contact, but not by airborne route and indirect contact.

**Figure 3.**
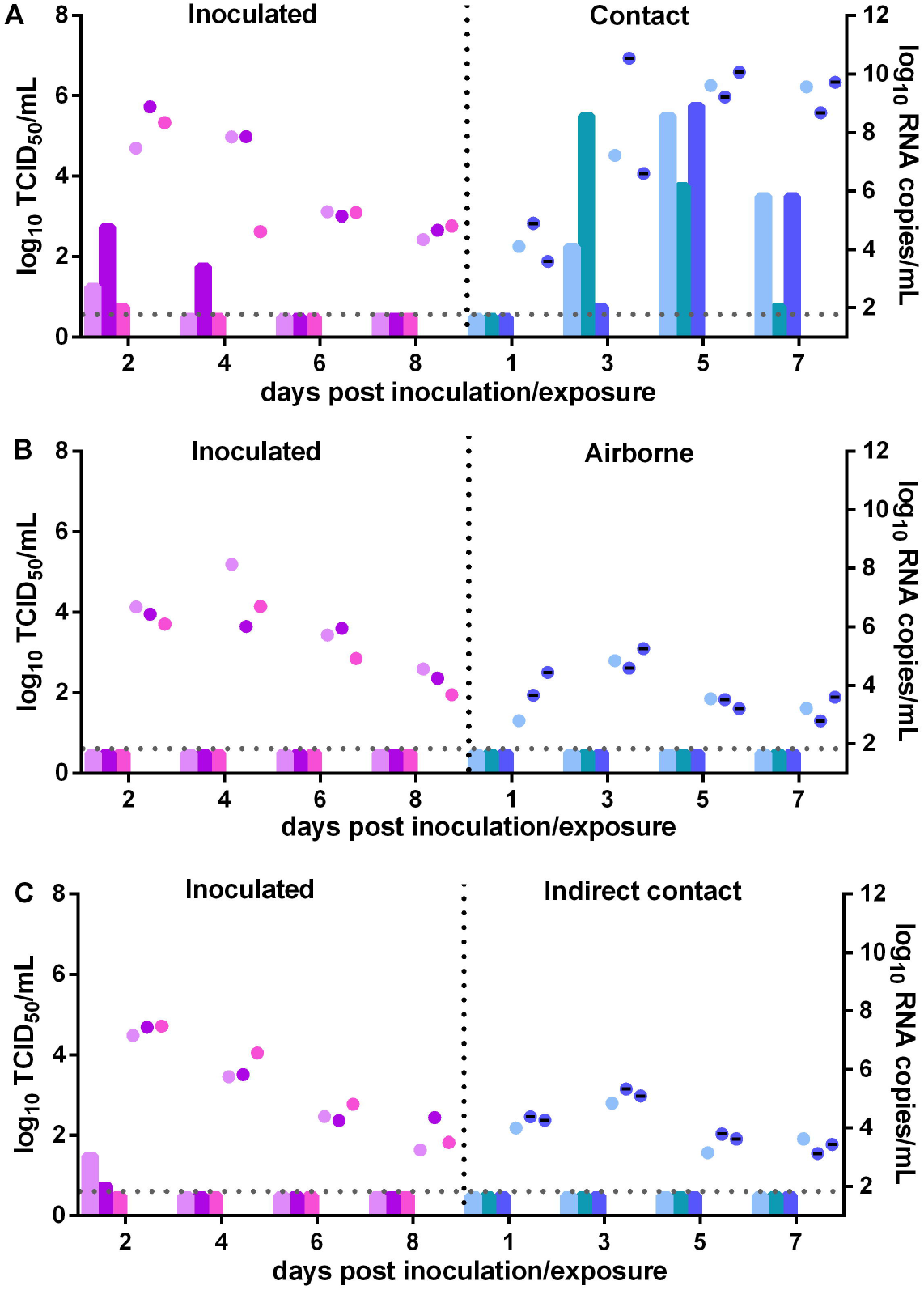
Transmission of SARS-CoV-2 from prior infected hamsters to naive hamsters. (A) Infectious viral load (log_10_TCID_50_ shown in bars) and viral RNA copies (log_10_RNA copies/mL, shown in dots with matched color) detected in nasal washes of prior infected hamsters inoculated with 10^6^ TCID_50_ of SARS-CoV-2 and the naïve contact hamsters. At 24 hours’ inoculation, the prior infected donor hamsters and the naïve contact hamsters were co-housed together in a new cage. (B) Viral titers and viral RNA copies detected in nasal washes of the prior infected hamsters inoculated with SARS-CoV-2 and naive hamsters in airborne transmission group. At 24 hours’ inoculation, the prior infected donor hamsters and naive hamsters were transferred to an airborne transmission cage. (C) Viral titers and viral RNA copies detected in nasal washes of prior infected hamsters inoculated with SARS-CoV-2 and naive hamsters in indirect contact transmission group. At 48 hours’ inoculation, the donor hamsters were removed and transferred to a new cage and the naive hamsters were placed into the cage housing the prior infected donor hamsters before. Nasal washes were collected from all hamsters every other day for viral titration and RNA quantification.

### Transmission of SARS-CoV-2 between prior infected hamsters

We evaluated the potential transmission of SARS-CoV-2 between prior infected hamsters by direct contact and airborne route. For direct contact transmission, four prior infected donor hamsters were inoculated with 10^6^ TCID_50_ of the virus. At two hours’ inoculation, another four prior infected hamsters were co-housed together with those four donors in a new cage. A high level of virus titer was detected in nasal washes of the donor hamsters at 1 and 3 dpi, averagely 10^3.56^ TCID_50_/mL and 10^2.25^ TCID_50_/mL, while viral RNA copies were maintained at 10^7^ to 10^8^copies/mL. Since 5 dpi, no live virus was found in nasal washes of the donors. At 3 dpe, live virus was detected in nasal washes in one contact hamster, with a titer of 10^4.25^ TCID_50_/mL, and viral RNA load in this hamster was also greatly improved to 10^8.26^copies/mL (Figure 4A). At 5 dpe, live virus was detected in nasal washes of another hamster in the contact transmission group, with a low titer of 10^0.75^ TCID_50_/mL, while viral RNA copies were improved to 10^6^copies/mL. At 7 dpe, virus titer in nasal washes of this hamster was substantially improved to 10^3.25^ TCID_50_/mL, and viral RNA copies were further improved to 10^8.46^copies/mL (Figure 4A). It seems that the virus was first transmitted from the artificially inoculated hamsters to a prior infected contact hamster, and then was sequentially transmitted to another prior infected contact hamster. The experiment was also performed similarly at 24 hours’ inoculation, four prior infected donors and four prior infected contact hamsters were co-housed in a new cage at 24 hours’ inoculation. No live virus was detected in nasal washes of all contact hamsters during the experiment period (Figure S5A). The experiment results showed that SARS-CoV-2 was transmitted between prior infected hamsters by direct contact during a very short period of early infection. For airborne transmission, three prior infected donor hamsters were inoculated with 10^6^ TCID_50_ of the virus and another three prior infected hamsters were as the recipients in airborne transmission group. The three donor hamsters and the three recipient hamsters were transferred to an airborne transmission cage at two hours’ or 24 hours’ inoculation, no live virus was detected in nasal washes of all recipient hamsters in the two airborne transmission groups (Figure 4B & Figure S5B). The results showed that SARS-CoV-2 was not transmitted between prior infected hamsters by airborne route. Taken together, SARS-CoV-2 has limited transmission ability between prior infected hamsters by direct contact during a short period of early infection, but without the ability to transmit by airborne route.

**Figure 4.**
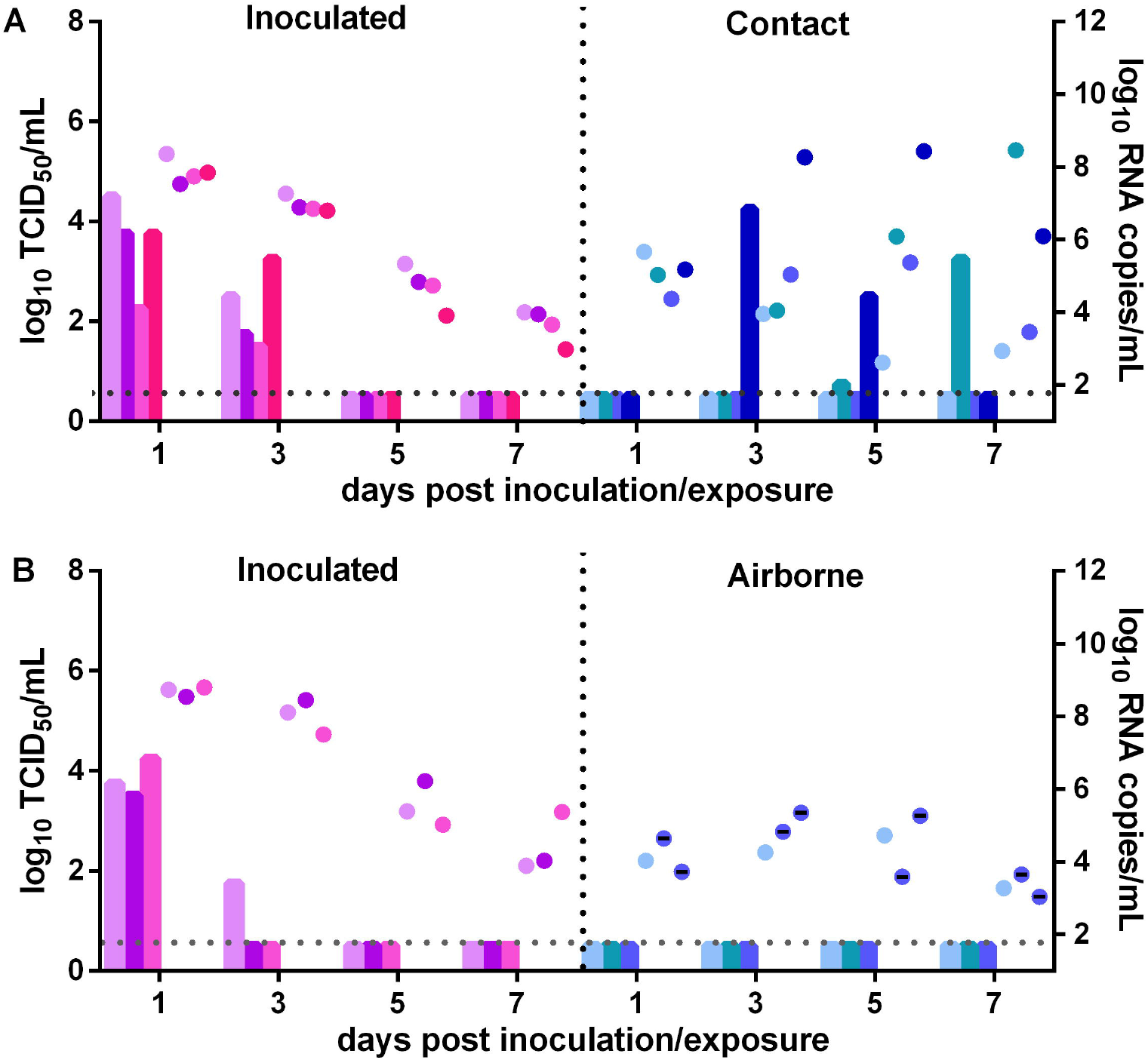
Transmission of SARS-CoV-2 between prior infected hamsters. A. Infectious viral load (log_10_TCID_50_ shown in bars) and viral RNA copies (log_10_RNA copies/mL, shown in dots with matched color) detected in nasal washes of prior infected hamsters inoculated with 10^6^ TCID_50_ of SARS-CoV-2 and the prior infected contact hamsters. At two hours’ inoculation, the donor hamsters and the contact hamsters were co-housed together in a new cage. (B) Viral titers and viral RNA copies detected in nasal washes of the prior infected donor hamsters inoculated with SARS-CoV-2 and the prior infected hamsters in airborne transmission group. At two hours’ inoculation, the donor hamsters and other hamsters were transferred to the airborne transmission cage. Nasal washes were collected from all hamsters every other day for viral titration and RNA quantification.

### Impact of a lower dose inoculation on SARS-CoV-2 transmission

We evaluated the impact of a lower dose inoculation on SARS-CoV-2 transmission between prior infected hamsters and naive hamsters by direct contact. For transmission of the virus from naive donors to prior infected contacts, four naive donor hamsters were inoculated with 10^4^ TCID_50_ of the virus, and at 24 hours’ inoculation, the donors and another four prior infected hamsters were co-housed together in a new cage. Similar to the high dose inoculation, viral load in nasal washes was maintained at a higher level at 2 and 4 dpi as well, and then followed a rapid decline (Figure 5A). Viral RNA copies were maintained at about 10^9^copies/mL during the first four infection days, and then slowly declined. Live virus was detected in two contact hamsters at 1 dpe, one with a very low titer of 10^0.75^ TCID_50_/mL. At 3 dpe, live virus was found in nasal washes of all four prior infected hamsters, two of which was significantly improved than before (Figure 5A). Viral RNA copies in nasal washes at 3 dpe were also significantly improved than before. Obviously, SARS-CoV-2 was efficiently transmitted from naive donor hamsters to prior infected hamsters by direct contact. For transmission of the virus from prior infected hamsters to naive hamsters, four prior infected donor hamsters were inoculated with 10^4^ TCID_50_ of the virus. At 24 hours’ inoculation, the donor hamsters and another four naive hamsters were co-housed together in a new cage. At 2 dpi, virus load in nasal washes of the donor hamsters was at a moderate level, about 10^2.25^TCID_50_/mL, and viral RNA copies were about 10^7.8^copies/mL (Figure 5B). Live virus was detected in nasal washes of three of the four naive contacts at 3 dpe, with titers from 10^3.25^ TCID_50_/mL to 10^5.5^ TCID_50_/m (Figure 5B). At 5 dpe, another contact hamster was also infected by SARS-CoV-2, with a high viral titer of 10^5.25^ TCID_50_/mL in nasal washes. Viral RNA copies were rapidly improved to 10^8.72^copies/mL at 3 dpe, and further improved to 10^10.15^copies/mL at 5 dpe, then slowly declined at 7 dpe. The results showed that SARS-CoV-2 was efficiently transmitted from prior infected donor hamsters to the naïve contacts. In summary, with a lower inoculation, SARS-CoV-2 was still efficiently transmitted between prior infected hamsters and naive hamsters.

**Figure 5.**
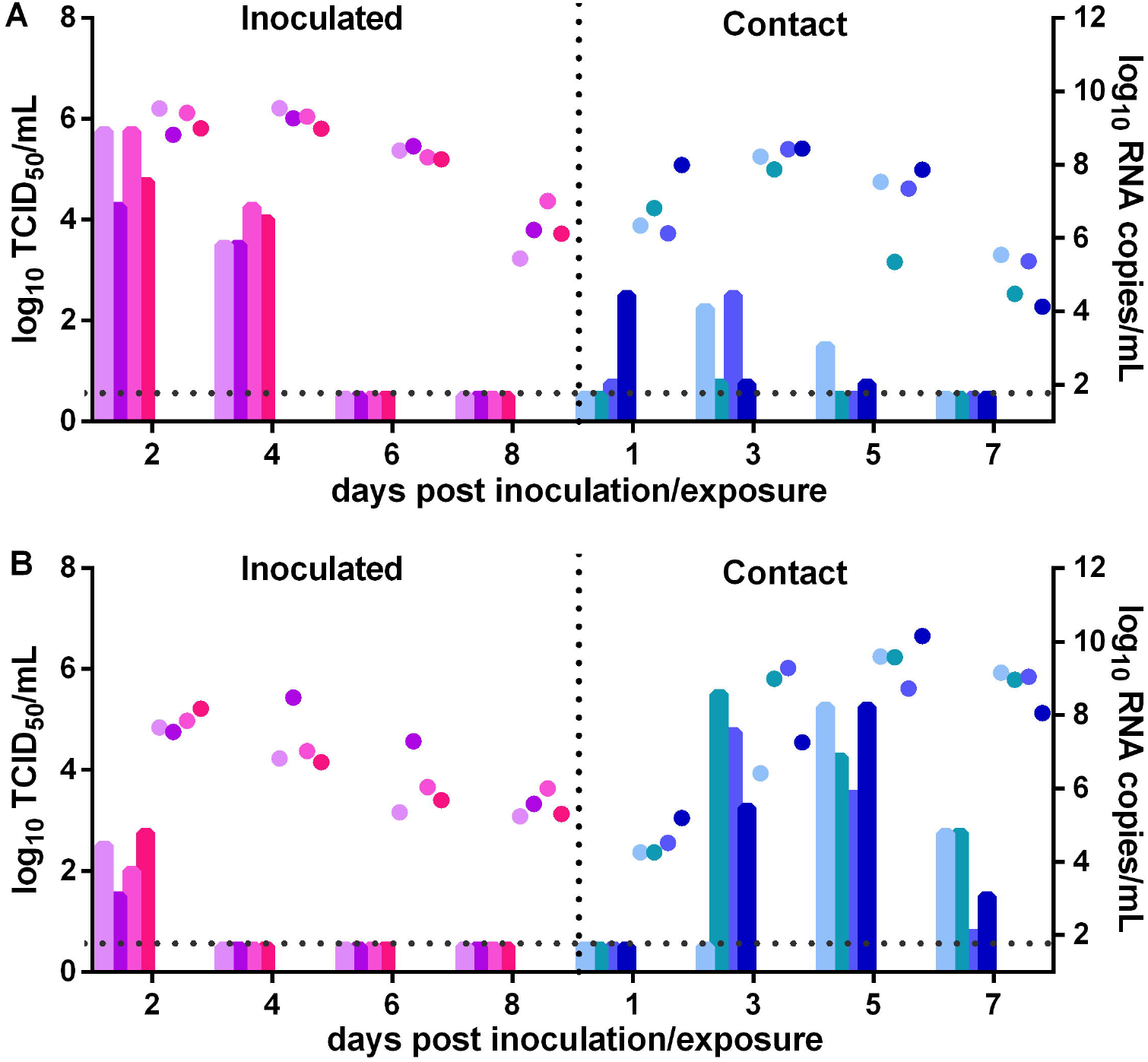
Impact of a lower dose infection on SARS-CoV-2 transmission between naïve hamsters and prior infected hamsters. (A) Infectious viral load (log_10_TCID_50_ shown in bars) and viral RNA copies (log_10_RNA copies/mL, shown in dots with matched color) detected in nasal washes of the naive hamsters inoculated with 10^4^ TCID_50_ of SARS-CoV-2 and the prior infected contact hamsters. At 24 hours’ inoculation, the naïve donor hamsters and the prior infected contact hamsters were co-housed together in a new cage. (B) Viral titers and viral RNA copies detected in nasal washes of the prior infected donor hamsters inoculated with 10^4^ TCID_50_ of SARS-CoV-2 and naive contact hamsters. At 24 hours’ inoculation, the prior infected donor hamsters and naïve contact hamsters were co-housed together in a new cage. Nasal washes were collected from all hamsters for viral titration and RNA quantification.

## Discussion

Our results showed that prior infection of SARS-CoV-2 elicited a higher titer of neutralizing antibodies in hamsters, and provided protective immunity against SARS-CoV-2 re-challenge. However, it was not sterizing immunity, and the virus still moderately replicated in nasal turbinates of prior infected hamsters, indicating that prior infected hamsters can be artificially re-infected after a short recovery period, even with a high level of neutralization antibodies. The conclusion is consistent with recent reports showing that recovered COVID-19 patients were re-infected in the presence of neutralizing antibodies^24,25^. A large study of a recovered cohort of 175 COVID-19 patients revealed that 6% of COVID-19 patients did not show any antibody response at all, and about 30% COVID-19 patients showed very low neutralizing antibodies^26^. Considering the gradual decay of neutralizing antibodies^27–30^ and a considerable population with very low neutralizing antibodies, the re-infection of some recovered COVID-19 patients will be unavoidable in the future. We also showed that prior infected hamsters can be naturally re-infected by direct contact or airborne route. The results of transmission study showed that SARS-CoV-2 can be transmitted effectively from naive hamsters to prior infected hamsters by direct contact and airborne routes, but not by indirect contact. Additionally, SARS-CoV-2 can be transmitted effectively from prior infected hamsters to naive hamsters by direct contact, but not by airborne route and indirect contact. Furthermore, SARS-CoV-2 can be transmitted between prior infected hamsters by direct contact during a very short period of early infection, but the transmission efficiency was limited. Taken together, prior infection substantially reduced the transmission efficiency of SARS-CoV-2 from prior infected hamsters to the naïve or prior infected hamsters by airborne route, but had limited impact on lowering the transmission by direct contact. In contrast with SARS-CoV-2, seasonal influenza A virus transmission between ferrets can be substantially reduced or blocked by natural infection or vaccination with live attenuated viruses^31,32^. The underlying mechanism behind the difference is not clear. Given the facts of re-infection, effective transmission between the prior infected hamsters and the naive, and waning immunity of the recovered COVID-19 patients^28,29^, it would be much more difficult to achieve herd immunity by natural infection or vaccination. A much higher vaccination coverage rate may be needed. At present, many governments and public health agencies are considering introducing immunity passport to help with recovery of social community and economy activities^33^, but evidence supporting this proposal is not enough. Reducing or blocking SARS-CoV-2 transmission is critical for COVID-19 pandemic control. A roaring increase in confirmed infections and hospitalized patients may lead to the collapse of the health care system, resulting in more deaths, social panic and even economic paralysis. How does vaccination with COVID-19 vaccines impact SARS-CoV-2 transmission in humans? There is still no clear-cut answer. Recent studies demonstrated that intranasal immunization with an Ad vector vaccine provided near complete sterizing immunity to SARS-CoV-2 in mice^34^, which might block the virus transmission in humans. Further studies to evaluate the blocking efficiency of different vaccines vaccinated by different routes, in particular, intranasal vaccination, on SARS-CoV-2 transmission in humans and animal models are urgently needed. Our work will help to determine the herd immunity threshold more accurately, make more reasonable public health decisions, as well as aid the implementation of appropriate public health and social measures to control COVID-19.

## Acknowledgements

This research was supported by the National Natural Science Foundation of China (32000134) and the National Major Research & Development Program (2020YFC0840800). We thank all staffs at Biosafety Level 3 Laboratories of Military Veterinary Research Institute for their all support and help.

## Author Contributions

CMZ, YWG conceived and designed the project, CMZ, CZ, ZDG, NL, HC, LNL, LZ, KYM and SSZ performed the experiments. CMZ, CZ, ZDG and NL analyzed the data. CMZ drafted the manuscript, YWG, CFQ, ZDG and JXL revised the manuscript critically.

## Data sharing

Data will be made available on request, directed to corresponding author CMZ.

