## supplemental Table 1-3 and supplementary figures 1-5 for "Impact of Prior Infection on Protection and Transmission of SARS-CoV-2 in Golden Hamsters"

^#^ Correspondence author:

**Supplementary Methods**

**Ethics and biosecurity statement**

All animal studies were conducted in strict accordance with the Guide for the Care and Use of Laboratory Animals of Military Veterinary Research Institute. All animal experiments were approved by the Animal Care and Use Committee of Military Veterinary Research Institute and were performed by certified staffs. All experiments involving the infectious SARS-CoV-2 virus were performed in Animal Biosafety Level-3 (ABSL-3) facilities of Military Veterinary Research Institute.

**Virus and cells**

BetaCoV/Beijing/IME-BJ05-2020 (abbreviated as V34) was isolated and propagated in Vero-E6 cells, and the virus stock was titrated into TCID_50_/mL by the classical 50% endpoint assay (TCID_50_ assay) based on virus-induced cytopathic effect (CPE) in Vero-E6 cells. The viral titer of the stock is 10^7.0^ TCID_50_/mL.Vero-E6 cells were grown in DMEM containing 10% fetus calf serum, 100 U / mL penicillin and100 μg/ml streptomycin.

**SARS-CoV-2 challenged studies in golden hamsters**

To evaluate the protective immunity of prior infection against SARS-CoV-2, 24 golden hamsters, about 4-5 weeks old, were randomized into two groups: high dose infected group (HD)and low dose infected group (LD). Hamsters in HD were anesthetized with isoflurane and intranasally inoculated with 10^5^ TCID_50_ of the SARS-CoV-2 virus in 100 μL DMEM, and hamsters in LD were intranasally inoculated with 10^3^ TCID_50_ of the SARS-CoV-2 virus in 100 μL DMEM. Four hamsters in each group were euthanized to collect serum samples at 21 days post infection. Other hamsters in HD and LD were re-challenged with 10^6^ TCID_50_ of theSARS-CoV-2 virus in 100 μL DMEM. As the infected control (IC), another six naive hamsters were inoculated with 10^6^ TCID_50_ of the SARS-CoV-2 virus in 100 μL DMEM. At 2 and 4 dpi, nasal washes were collected from each hamster using 1 mL PBS. Half of the hamsters in each group were euthanized to obtain nasal turbinates and lungs following the nasal washes, one part was homogenized into 1 mL PBS, and another part was fixed in 2.5% glutaraldehyde. All collected nasal washes and the supernatants of the homogenized nasal turbinates and lungs were used for virus titration in Vero-E6 cells and for virus RNA quantification using real-time RT-qPCR. The glutaraldehyde-fixed nasal turbinates were used for transmission electron microscopy examination. The nasal turbinates and lungs were lysed with the buffer RLCK for 10 minutes and then for subgenomic mRNA quantification.

**SARS-CoV-2 transmission studies in golden hamsters**

As described in the part of SARS-CoV-2 challenged studies in golden hamsters, hamsters were also divided into two groups: HD and LD, and intranasally inoculated with 10^5^ TCID_50_ of the SARS-CoV-2 virus and 10^3^ TCID_50_ of the SARS-CoV-2 virus respectively. At 21 days post infection, these hamsters were used for SARS-CoV-2 transmission studies. The animal composition of each experiment group was shown in table S2 and table S3. The transmission routes of the SARS-CoV-2 virus include direct contact transmission, indirect contact transmission and airborne transmission. For direct contact transmission evaluation, three or four donor hamsters were anesthetized with isoflurane and intranasally inoculated with10^6^ TCID_50_ of the SARS-CoV-2 virus, and housed in a negative pressure independent ventilation cage. Two hours or twenty-four hours after inoculation, another three or four recipient hamsters in the transmission group were transferred to a new cage and co-housed together with the donor hamsters (Figure S2A). At 1, 3, 5 and 7 days post exposure (dpe), nasal washes were collected from the donor hamsters and the recipient hamsters. For airborne transmission evaluation, three donor hamsters were inoculated with 10^6^ TCID_50_ of the SARS-CoV-2 virus and housed in a separate cage. Two hours or twenty-four hours after inoculation, the three donor hamsters and another three recipient hamsters were transferred to a new airborne transmission cage (Figure S2B), with two-wire-mesh partitions that prevented direct and indirect contact between animals and allowed the spread of SARS-CoV-2 via the flow air. The distance between the two-wire-mesh partitions was 2cm. Air flows from the donor cage to the cage housing recipient hamsters. Nasal washes were collected from the donor hamsters and those recipient hamsters in the transmission group at 1, 3, 5 and 7 dpe. For indirect contact transmission evaluation, three donor hamsters were inoculated with 10^6^ TCID_50_ of the SARS-CoV-2 virus and housed in a new cage. After 48 hours’ inoculation, the donor hamsters were removed and transferred to another new cage, and another three recipient hamsters were placed into the initial cage housing the donor hamsters. Nasal washes from the recipient hamsters were collected at 1, 3, 5 and 7 dpe, while nasal washes from donor hamsters were collected at 2, 4, 6 and 8 dpi. The infectious virus loads in these samples were determined by the classical TCID_50_ assay and virus RNA copies were quantified using real-time qPCR. For transmission of the virus from naive hamsters to prior infected hamsters, the naive hamsters were as the donors and prior infected hamster were as the recipients. Inversely, for transmission of the virus from prior infected hamsters to the naive hamsters, prior infected hamsters were as the donors and the naive hamsters were as the recipients. For transmission of the virus between prior infected hamsters, randomly selected prior infected hamsters were as the donors and the recipients. For the lower dose infection on SARS-CoV-2 transmission, the donor hamsters were anesthetized with isoflurane and intranasally inoculated 10^4^ TCID_50_ of the SARS-CoV-2 virus, and then treated similarly as hamsters in other transmission experiments.

**Virus RNA quantification by quantitative real-time RT-PCR**

RNA was extracted from 200 μL samples using the viral RNA minikits (QIQGEN, Hilden, Germany) according to the manufacture’s protocol and eluted with 90 μL water, and 15μL RNA was used for the real-time qPCR to detect the N gene of SARS-CoV-2 using the Detection Kits for 2019-Novel Coranavirus RNA (Shenzhen Puruikang Biotech, China). The experiments were performed with an ABI7500 system (Roche, Switzerland). The amplification reaction conditions were 50℃ for 20 min for reverse transcriptase, followed by 95℃ for 3 min, and then 45 cycles of 95℃ for 5 s, 57℃ for 45 s, and finally 25℃ for 10 min. The SARS-CoV-2 RNA copies in each samples was estimated from the measured cycle threshold (Ct) values based on the established standard curves with a standard plasmid of the SARS-CoV-2 virus.

**Virus subgenomic RNA quantification**

The subgenomic RNA (sgmRNA) of the SARS-CoV-2 virus was a better indicator of virus replication in vivo. To measure virus replication in golden hamsters, the sgmRNA copies in nasal turbinates and lungs from the challenged hamsters were determined. The nasal turbinates and lungs were homogenized and lysed in 1 mL of the buffer RLCK (QIQGEN, Hilden, Germany) for 10 min, and centrifuged at high speed of 10000 rpm for 5 min. RNA was extracted from 500 μL of the centrifuged supernatants and eluted with 90 μL water. A pair of primers and a Taqman probe was designed targeting the E gene sgmRNA of SARS-CoV-2^1^. 9 μL RNA was used for real-time qPCR to detect the E gene sgmRNA using the One Step PrimeScriptTM III RT-qPCR Mix (Code No: RR600A, Takara) in an ABI7500 system (Roche, Switzerland). The reaction conditions were 52℃ for 5 min for reverse transcriptase, followed by 95℃ for 10 s, and then 40 cycles of 95℃ for 5 s, 60℃ for 30 s. The E gene sgmRNA copies in each samples was estimated from the measured cycle threshold (Ct) values based on a fitted standard curve with a serial 10-fold dilutions of a standard plasmid of the SARS-CoV-2 virus.

Infectious viral load determination by the classical TCID_50_ assay

Serial ten-fold dilutions of nasal washes and the supernatants of homogenized tissues with 1 mL PBS were added to Vero-E6 cells with 80% confluence in 96-well plates, and incubated for another 4 days at 37 ℃ with 5% CO2. The cytopathic effect was observed under an inverted microscopy. The viral titers were calculated using the Reed-Muench method

**Virus neutralization assay**

One hundred micro liters of virus dilutions, containing 100 TCID_50_of the SARS-CoV-2 virus, was incubated with 100 μL two-fold serial dilutions of the infected hamster serum or mock serum for 1 h, and200 μL of the mixtures were added to Vero-E6 cells with 80% confluence in 96-well plates, and incubated for another 4 days at 37 ℃with 5% CO2. The cytopathic effects were observed under a microscopy in the Animal Biological Safety Level-3 laboratories. Each sample has three replicates. The virus neutralizing antibody titers (NT_100_) were determined as the reciprocal number of the highest serum dilution that completely prevented the cytopathic effect.

**Transmission electron microscopy**

Nasal tissues were fixed in phosphate buffered 2.5% glutaraldehyde for 24 h. Specimens were then postfixed in 1% osmium tetroxide, washed, dehydrated with a series of ethanol and alcohol gradient, and embeded in a mixture of epoxy resin. The 70 nm-thick sections were stained with uranyl acetate and lead citrate, and examined in a 120 KV transmission electron microscopy by a special electron microscopy technician.

**Statistics analysis**

The one-way analysis of variance (ANOVA) and Tukey’s multiple comparisons test were used to analyze the statistical differences of viral titers, RNA copies and sgmRNA copies in nasal washes, nasal turbinates and lungs between different experimental groups at 2 and 4 dpi. Mann-Whitney U test was used to analyze the difference of serum neutralizing antibody titers between HD and LD (p > 0.05, not significant, [ns]; p < 0.05, *；p <0.01, **; p < 0.001, ***). All data was analyzed with the software GraphPad Prism 6.02.

Supplementary tables

Table S1 grouping and treatments of animal experiments

| Grouping | n | Experimental treatment | |
| --- | --- | --- | --- |
|  |  | first infection dose | re-challenge dose |
| Infected Control | 6 | no | 10^6^ |
| HD | 8 | 10^5^ | 10^6^ |
| LD | 8 | 10^3^ | 10^6^ |

HD: high dose infected.

LD: low dose infected.

Table S2 animal constitutes of the transmission experimental groups.

| Graph ID | Transmission  routes | The donor group | | The recipient group | |
| --- | --- | --- | --- | --- | --- |
|  |  | n | constitution | n | constitution |
| Figure2A | DC | 3 | 3N | 3 | 2H+1L |
| Figure2B | AR | 3 | 3N | 3 | 1H+2L |
| Figure2C | IDC | 3 | 3N | 3 | 2H+1L |
| Figure3A | DC | 3 | 2H+1L | 3 | 3N |
| Figure3B | AR | 3 | 1H+2L | 3 | 3N |
| Figure3C | IDC | 3 | 2H+1L | 3 | 3N |
| Figure4A | DC | 4 | 2H+2L | 4 | 2H+2L |
| Figure4B | AR | 3 | 2H+1L | 3 | 2H+1L |
| Figure5A | DC | 4 | 4N | 4 | 2H+2L |
| Figure5B | DC | 4 | 2H+2L | 4 | 4N |

H: animals initially infected with a high dose of 10^5^ TCID_50_ of SARS-CoV-2.

L: animals initially infected with a lower dose of 10^3^ TCID_50_ of SARS-CoV-2.

N: naive animals

DC: direct contact transmission

AR: airborne transmission

IDC: indirect contact transmission

Table S3 animal constitutes of the transmission experimental groups in the supplementary

| Graph ID | Transmission  routes | The donor group | | The recipient group | |
| --- | --- | --- | --- | --- | --- |
|  |  | n | constitution | n | constitution |
| Figure S4A | DC | 3 | 1H2L | 3 | 3N |
| Figure S4B | AR | 3 | 1H2L | 3 | 3N |
| Figure S5A | DC | 4 | 2H+2L | 4 | 2H+2L |
| Figure S5B | AR | 3 | 2H+1L | 3 | 2H+1L |

*The meaning of H, L, N, DC and AR are the same as that explained in Table S2.

Supplementary figure legends

**Figure S1** The neutralizing antibodies against SARS-CoV-2 in serum samples of hamsters in HD and LD at 21 dpi. The statistic difference between HD and LD was analyzed by Mann-Whitney U test (p > 0.05, not significant, [ns]; p < 0.05, *).

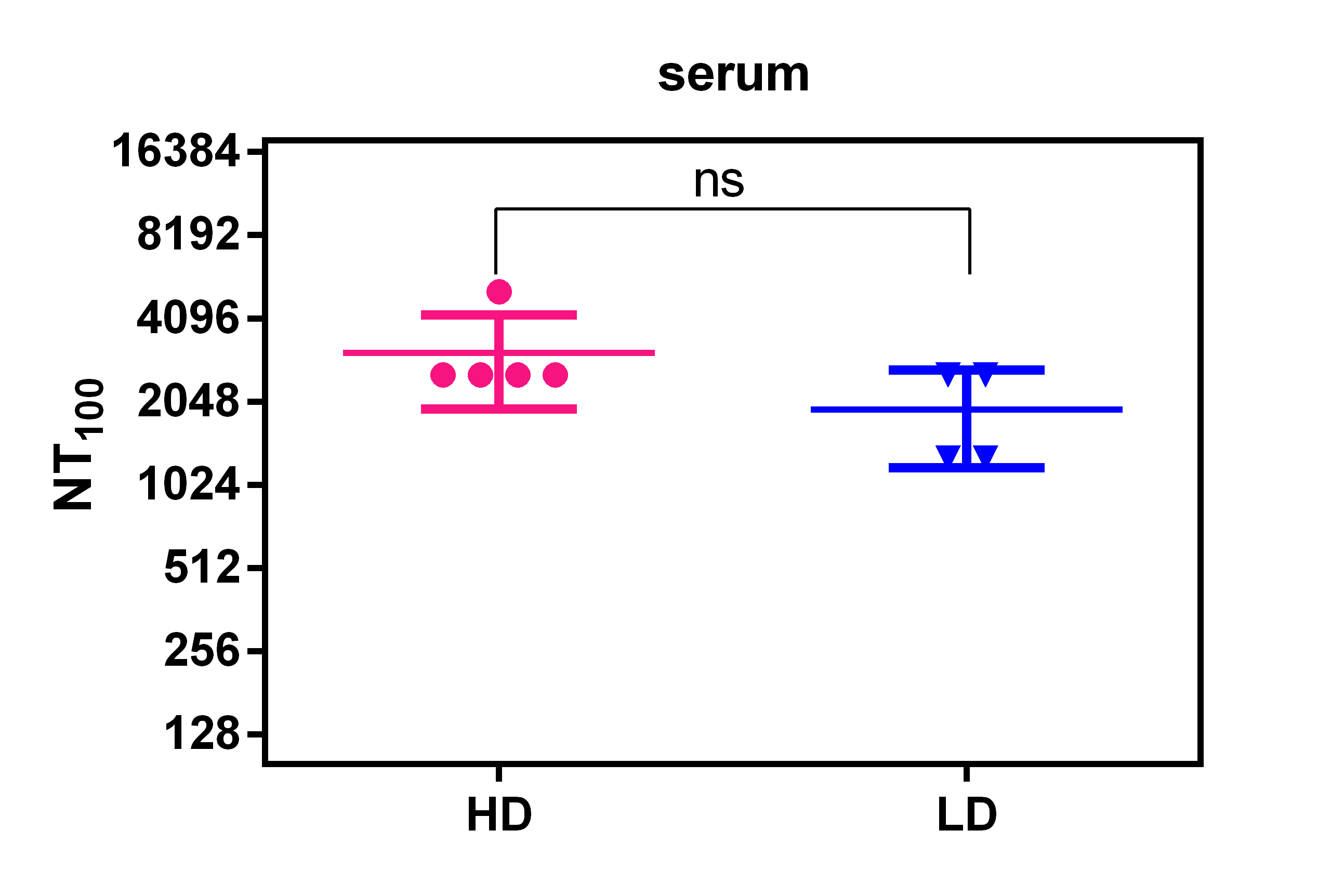

**Figure S2** Electron microscopy examination for the presence of SARS-CoV-2 in nasal tissues of hamsters that was re-challenged with the virus. Nasal tissues were fixed in glutaraldehyde, embeded in epoxy resin, cut into 70 nm-thick sections, and stained with uranyl acetate and lead citrate for examination withtransmission electron microscopy. The coronavirus-like particles were labeled as red circles.

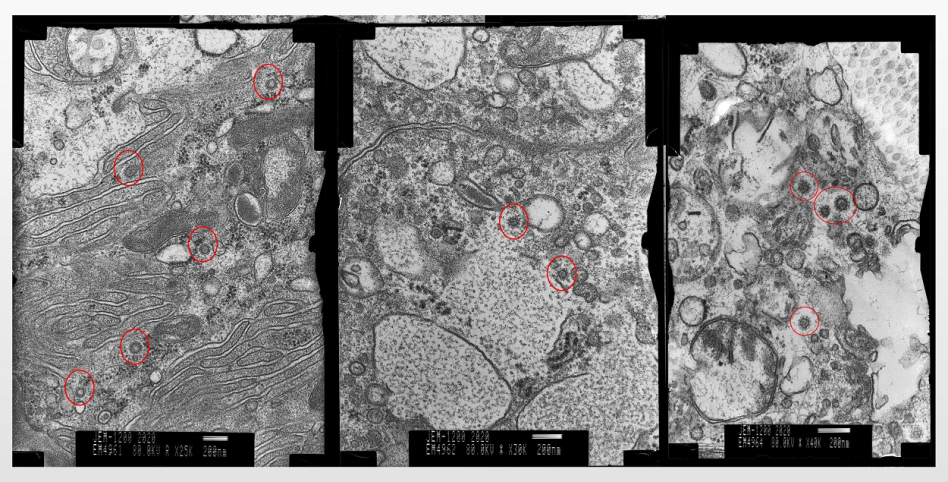

**Figure S3** Schematic representation of the transmission cages used for contact transmission studies (A) and airborne transmission studies (B)**.** The cage for airborne transmission studies has two wire mesh separators that prevent direct contact and indirect contact between animals and allow the spread of the SARS-CoV-2 virus through air flow.

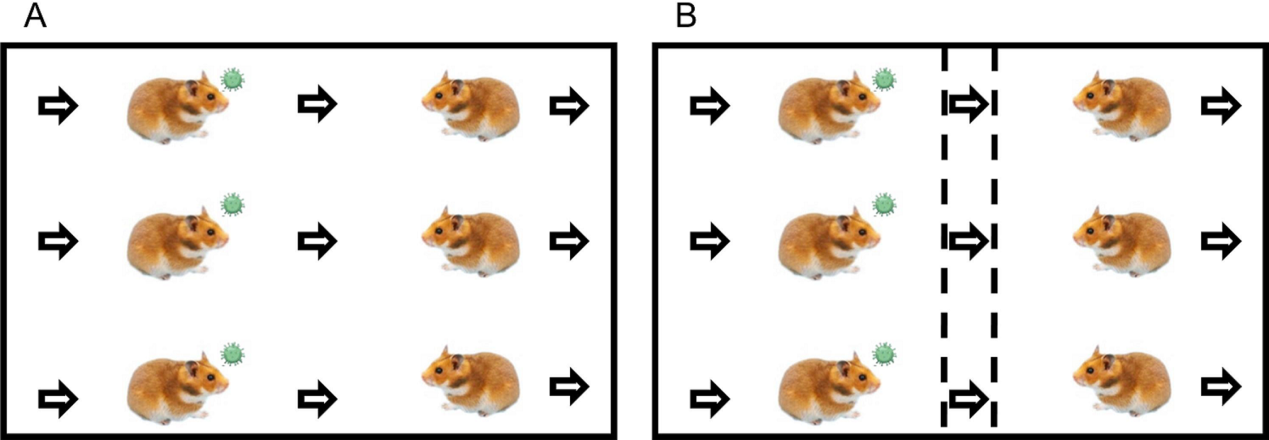

**Figure S4** Transmission of SARS-CoV-2 from prior infected hamsters to naive hamsters. (A) Infectious viral load (log_10_TCID_50_ shown in bars) and viral RNA copies (log_10_RNA copies/mL, shown in dots with matched color) detected in nasal washes of prior infected donor hamsters inoculated with 10^6^ TCID_50_ of SARS-CoV-2 and the naïve contact hamsters. At two hours’ inoculation, the donor hamsters and the contact hamsters were co-housed together in a new cage. (B) Viral titers and viral RNA copies detected in nasal washes of the prior infected donor hamsters inoculated with SARS-CoV-2 and naive hamsters in airborne transmission group. At two hours’ inoculation, the donor hamsters and the naïve recipient hamsters were transferred to an airborne transmission cage. Nasal washes were collected every other day for viral load and RNA quantification.

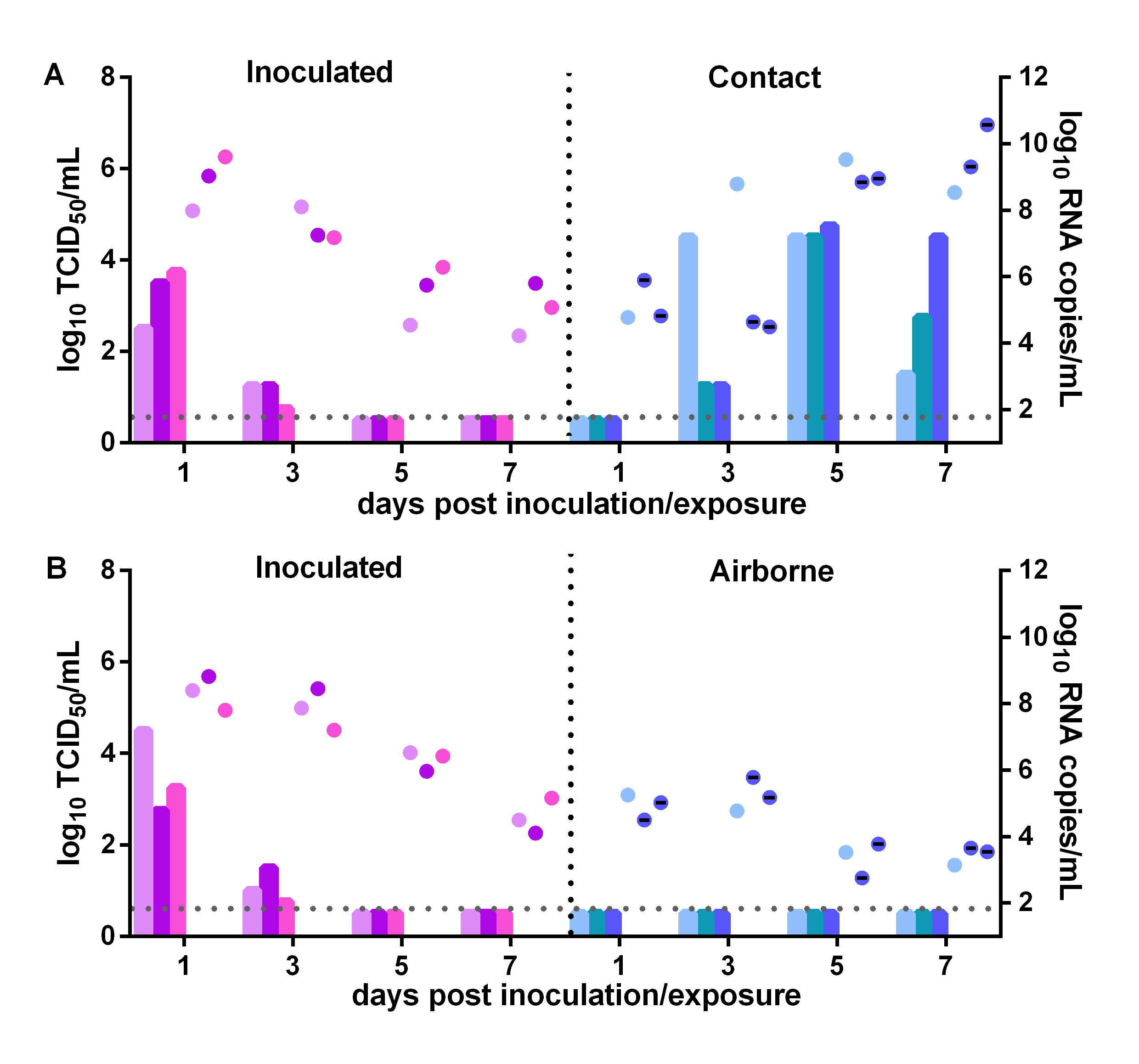

**Figure S5** Transmission of SARS-CoV-2 between prior infected hamsters. (A) Infectious viral load (log_10_TCID_50_ shown in bars) and viral RNA copies (log_10_RNA copies/mL, shown in dots with matched color) detected in nasal washes of prior infected donor hamsters inoculated with 10^6^ TCID_50_ of SARS-CoV-2 and the prior infected contact hamsters. At 24 hours’ inoculation, the prior infected donor hamsters and the prior infected contact hamsters were co-housed together in a new cage. (B) Viral titers and viral RNA copies detected in nasal washes of the prior infected donor hamsters inoculated with SARS-CoV-2 and the prior infected recipient hamsters in airborne transmission group. At 24 hours’ inoculation, the donor hamsters and the recipient hamsters were transferred to an airborne transmission cage. Nasal washes were collected from all hamsters every other day for viral load and RNA quantification.

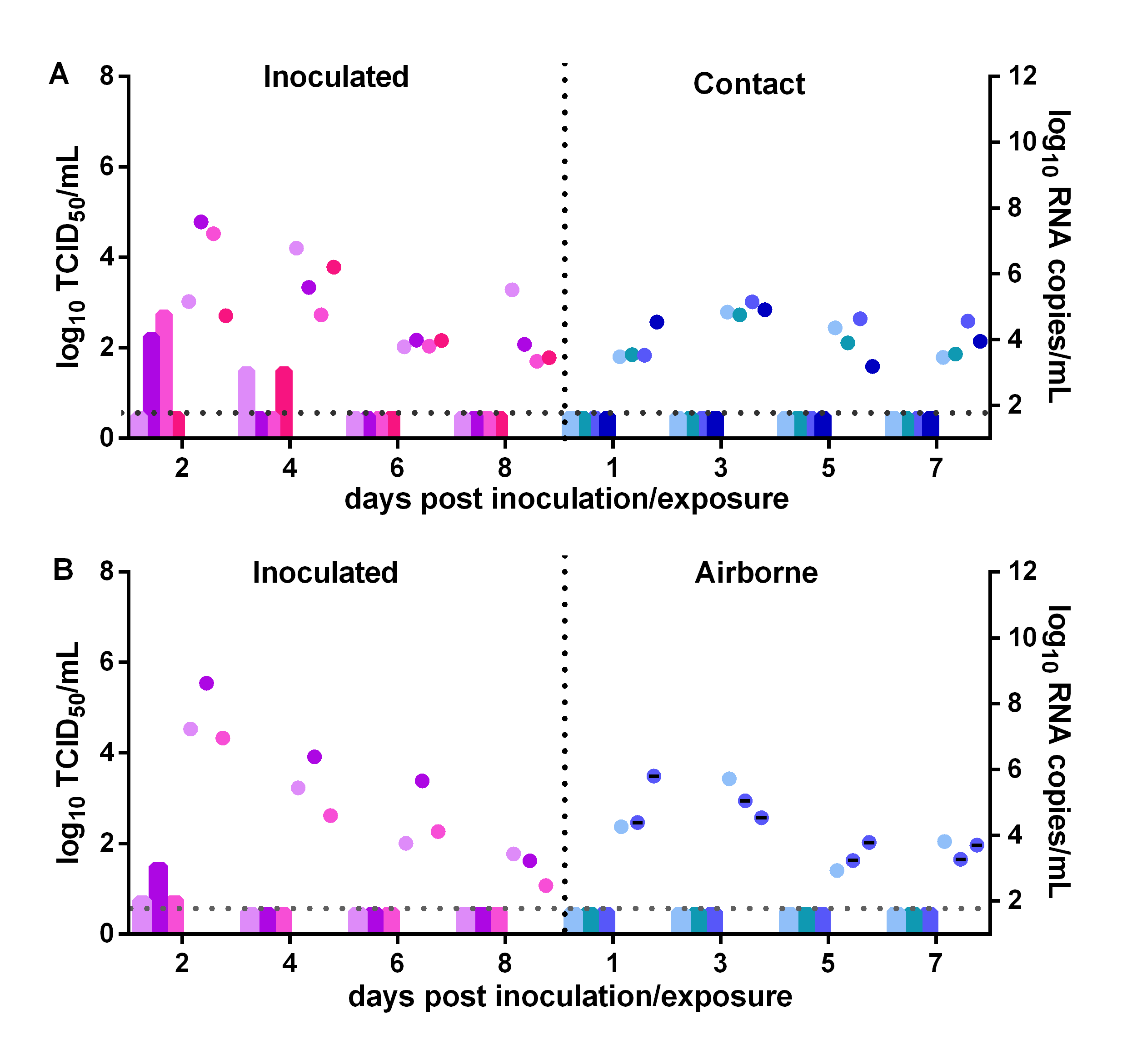
